# Climate change and chill accumulation: implications for tree fruit production in cold-winter regions

**DOI:** 10.1101/2020.08.26.268979

**Authors:** Hossein Noorazar, Lee Kalcsits, Vincent P. Jones, Matthew S. Jones, Kirti Rajagopalan

## Abstract

Winter chill accumulation is critical for the productivity and profitability of perennial tree fruit systems. Several studies have quantified the impacts of global warming on chill accumulation in the warmer production regions of the world, where insufficient chill events occur and their frequency is increasing. In contrast, we focus on a region with relatively cold winters–the Pacific Northwest United States (PNW)–where insufficient chill events are currently absent, and quantify the potential for introduction of these risks under climate change. Results identified spatial variation within the PNW, with chill accumulation projected to increase in some areas but decrease in others. There was also spatiotemporal variation in the driving factors of changes to chill accumulation. Even with decreases in chill accumulations, there are likely minimal issues with insufficient chill accumulation. However, delayed chill accumulation in combination with advances in the onset of heat accumulation can potentially shift the region from one where spring phenology is primarily forcing-driven to one where the dynamic interplay between chilling and forcing processes become important. These interactions might create production risks for varieties with high chill requirements, post mid-21^st^-century under high emissions scenarios. Future work should focus on understanding, modeling, and projecting responses across these overlapping chilling and forcing processes. Additionally, given significant spatial differences across a relatively small geographic range, it is also critical to understand and model these dynamics at a local landscape resolution for regions such as the PNW.

## 1 Introduction

Winter chill accumulation is critical for productive and profitable temperate tree-fruit production systems. Specifically, chilling affects the emergence from endodormancy, transition to ecodormancy, and the resumption of growth in spring (Saure, 1985). Commercial varieties have a breadth of chilling requirements that make them suitable for diverse regions (Darbyshire et al., 2016; Luedeling et al., 2011; Samish and Lavee, 1962). Insufficient chill accumulation has adverse impacts (e.g. increased flower bud abscission, non-uniform bud break, reduced flower and fruit quality), affecting both fruit and vegetative development (Atkinson et al., 2012). Currently, these issues are important in warmer temperate tree-fruit growing regions such as Israel, South Africa, Spain, and California where varietal selection can be often be based largely on chill requirements (Alburquerque et al., 2008; Baldocchi and Wong, 2008; Luedeling, Zhang and Girvetz, 2009; Luedeling, Zhang, Luedeling and Girvetz, 2009; Luedeling, Zhang, McGranahan and Leslie, 2009; Ruiz et al., 2007). While climate change impact studies on insufficient chill accumulation have focused on these currently at-risk areas (Baldocchi and Wong, 2008; Luedeling, Zhang and Girvetz, 2009; Luedeling, Zhang, Luedeling and Girvetz, 2009; Farag et al., 2010; Midgley and Lötze, 2008; Wrege et al., 2010; Parker and Abatzoglou, 2019; Kaufmann and Blanke, 2014), future warming and associated decreases or delays in winter chill accumulation can introduce risks in regions with historically cooler winters, and there are limited studies (Houston et al., 2018; Luedeling and Brown, 2011) focusing on such regions.

Dormancy in tree fruit species is a time period where buds remain inactive. Establishing more accurate terminology, Lang et al. (Lang et al., 1987) proposed two dormancy stages– endodormancy and ecodormancy– defined by physiological and environmental inhibitions of regrowth, respectively. The stages are not discrete, but dynamic in response to environmental stimuli (Fuchigami et al., 1982). Dormancy induction in fall is regulated by either photoperiod, or temperature, or a combination of both (Heide, 2008; Kalcsits et al., 2009; Tanino et al., 2010). Once dormancy is induced, a tree will remain endodormant until it fulfills its chilling requirement at which point it enters ecodormancy when environmental conditions prevent budbreak (e.g. temperatures are not warm enough); chilling and heating (forcing) requirements both need to be fulfilled for budbreak. The chill and heat accumulation periods can overlap (Yang et al., 2020). However, in relatively cool temperate climates, chill requirements are frequently met during early winter and spring phenology is primarily affected by the forcing requirements and minimally affected by chill accumulation (Guo et al., 2014; Yang et al., 2020). In contrast, in relatively warmer regions, which currently experience higher risks of insufficient chill accumulation, there can be high overlaps between chilling and forcing processes and the dynamic interplay between them are more relevant for spring phenology and bud break (Guo et al., 2014). With climate change, regions with cold winters can face delayed chill accumulation and advanced heat accumulation resulting in a regime shift: spring phenology shifts from one that is primarily forcing driven to one where the interplay between chilling and forcing processes become more important.

Our objective was to focus on a region with cooler winters - that does not currently face risks of insufficient chill accumulation - and understand the potential evolving risks under climate change. We take the US Pacific Northwest (PNW) as a case study. Although temperate tree fruit crops are grown throughout the continental US, the PNW is the largest apple and sweet cherry production region in the country. The PNW accounted for 67% of the US apple production in 2018 with a value of $2.1 billion, and 87% of the US fresh cherry production valued at $470 million (USDA NASS 2018 statistics). Cool wet winters and warm dry summers of the PNW are ideal for specialty crop production. Irrigation water availability and proximity to processors and markets provide an additional competitive advantage (Houston et al., 2018). Therefore, while risk of insufficient chill accumulation is currently insignificant in the PNW, given its position as the largest production area in the US for multiple tree fruit, it is an ideal case study to understand future production risks under warming.

Only two studies have addressed climate change impacts on chill accumulation for tree fruit in the PNW and they were either restricted to two locations with only one time frame (2020–2049) (Houston et al., 2018) or in the larger context of tree fruit production worldwide (Luedeling et al., 2011), and diverge in results. In the first study (Houston et al., 2018), high chill-requirement cultivars in one location faced insufficient chill accumulation as early as the 2030s, but was relatively unaffected in another. In contrast, the second study, which highlighted a location in Canada close to the northern part of the PNW, reported an increase in chill accumulation (Luedeling et al., 2011). Neither considered chill accumulation in the context of heat accumulation nor described potential shifts in processes that drive spring phenology. Unrelated to tree-fruit, Harrington and Gould (Harrington and Gould, 2015) noted potential reductions in chill accumulation in boreal tree species in the PNW.

A variety of temperature-based empirical models have been developed to estimate chill accumulation and the transition from endo-to ecodormancy. These include the Chilling Hour Model (Bennett et al., 1950), the Utah Model (Richardson et al., 1974), and the Dynamic Model (Erez and Fishman, 1997; Erez et al., 1989). The Dynamic Model takes into account the positive effect of cool temperatures, the negative effect of high temperatures, the positive effect of moderate temperatures, the effect of moderate temperatures alternating with chilling temperatures, and the time heterogeneity of chilling negation. The Dynamic Model has been reported to better capture the dynamic nature of the chilling process compared to other models (Darbyshire et al., 2016; Luedeling, 2012; Luedeling and Brown, 2011; Linsley-Noakes and Allan, 1994). This model has been reported to closely align with observations for a variety of crops and cultivars (Luedeling, Zhang and Girvetz, 2009; Miranda et al., 2013; Zhang and Taylor, 2011), and for a variety of climatic regimes (Luedeling, 2012; Pérez et al., 2008). Moreover, the Dynamic Model has been shown to work well in warmer climatic regimes (Luedeling, Zhang and Girvetz, 2009; Erez and Fishman, 1997; Luedeling, 2012)-critical in the context of climate change and a shift to warmer temperature regimes.

Our study takes a spatially-explicit approach that covers all current tree fruit production areas in the PNW and aims to (a) quantify changes to risk of insufficient chill accumulation as a result of climate change in colder tree-fruit production regions, and (b) describe factors that lead to this change in risk. We do this by driving the Dynamic Chilling Portions Model (Erez and Fishman, 1997; Erez et al., 1989) with historical climate and future climate projections, and accounting for shifts in the timing of conditions conducive to heat accumulation. The results of this study will improve our understanding of productions risks related to chill accumulation in regions that currently have relatively cooler winters. It can also help PNW growers develop adaptation strategies for potential new insufficient chill accumulation risks they are not currently exposed to, and sustain the region’s standing as the major apple and cherry production region in the US.

## 2 Methods

### 2.1 Study Area

Apple and cherry production areas in the PNW are primarily in Washington State with some production in Idaho and Oregon states. We identified tree fruit growing areas by overlaying two 2018 agricultural land use datasets on our gridded climate dataset (described below) and resulted in ∼2300 simulation grids (Fig. 1). For the Washington State part of our study area, we used the Agricultural Land Use Database created by the Washington State Department of Agriculture (WSDA, n.d.). For the remaining areas we used Cropland Data Layer created by the United States Department of Agriculture (USDA CDL, n.d.). To highlight the spread in results, we also picked four representative locations spanning the breadth of our study area (Fig. 1). Our locations spanned an elevation between 173.9 and 333.2 m.a.s.l., winter mean temperature between 4.2°C in Omak and 8.5°C in Eugene, and annual mean precipitation of 220.1 mm in Yakima and 1164.4 mm in Eugene (Table 1).

**Table 1.** Elevation and climate information for the four representative locations.

| Location | Elevation (m) | Ann. Temp. Mean (°C) | Winter Temp. Mean (°C) | Ann. Precip. Mean (mm) | Winter Precip. Mean (mm) |
| --- | --- | --- | --- | --- | --- |
| Omak | 318.7 | 9.9 | 4.2 | 312.1 | 203.8 |
| Yakima | 333.2 | 10.6 | 5.6 | 220.1 | 160.7 |
| Walla Walla | 272.9 | 12.2 | 7.4 | 459.7 | 317.7 |
| Eugene | 173.9 | 11.5 | 8.5 | 1164.4 | 934.2 |

**Fig. 1.**
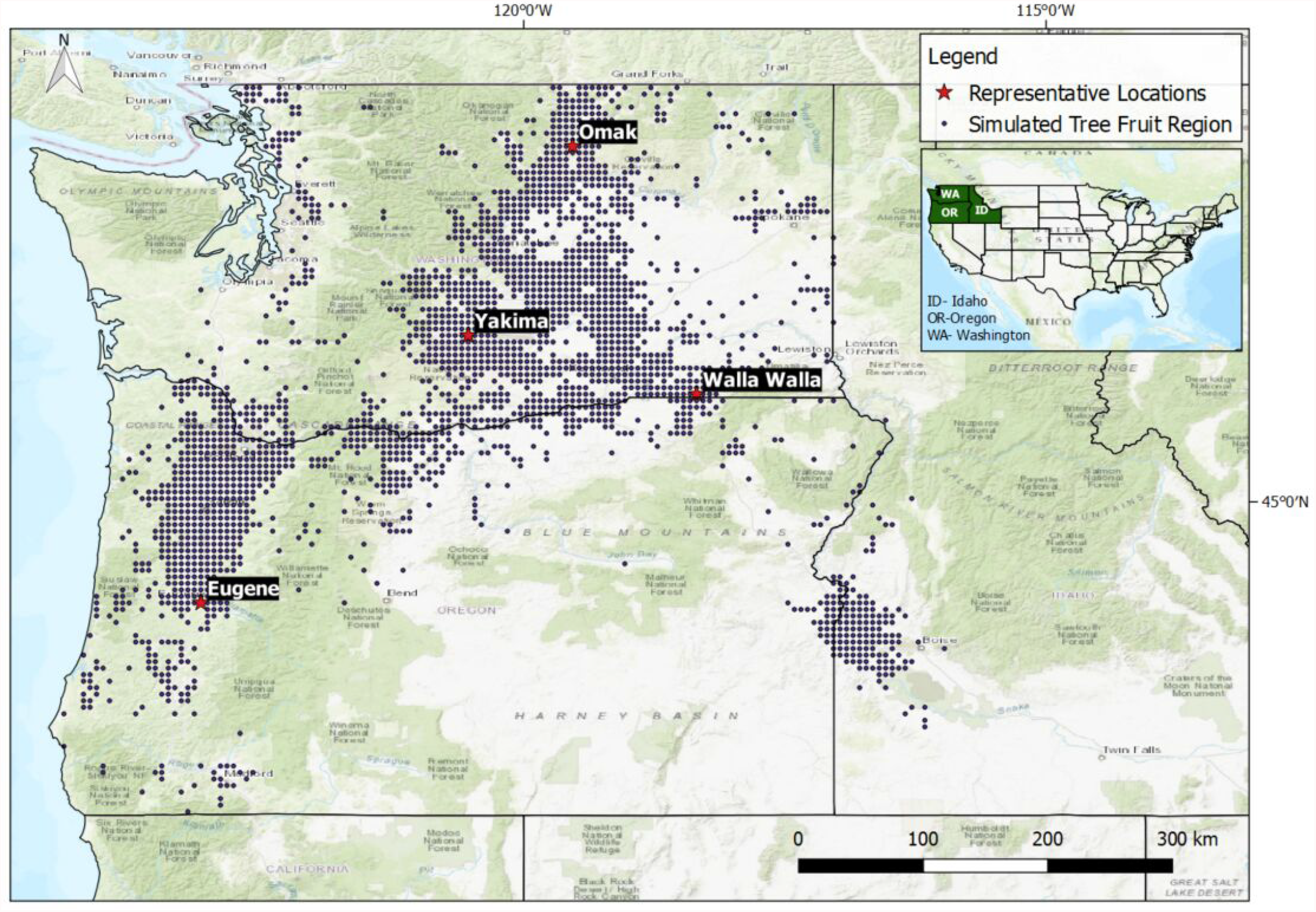
Map of the study domain. Black dots correspond to tree fruit growing areas that we simulated. Red stars correspond to four locations (Omak, Yakima, Walla Walla, Eugene) across the study domain that we selected to highlight some results in depth.

### 2.2 Historical Climate and Future Climate Projections

Historical simulations (1979–2016) were based on the gridded meteorological observations product -gridMET (Abatzoglou, 2013). Future climate projections (2006-2099) were based on (Abatzoglou and Brown, 2012), which downscaled and bias-corrected the Coupled Model Intercomparison Project 5 (CMIP5; (Taylor et al., 2012)) data to a 1/24th degree (4 km) spatial resolution based on the Modified Multivariate Adaptive Constructed Analog method (Abatzoglou and Brown, 2012). This data was subsequently re-gridded to a 1/16 degree (6km) resolution by linear interpolation for computational efficiency.

While the future simulations start from 2006, we use data starting from 2026 for aggregation into three time frames: 2026-2050, 2051-2075, and 2076-2099. Nineteen climate models under two representative concentration pathways (RCPs) were used for a total of 38 projections that captured the range in uncertainty in projections. The 19 models used in this study were bcc-csm1-1, bcc-csm1-1-m, BNU-ESM, CanESM2, CCSM4, CNRM-CM5, CSIRO-Mk3-6-0, GFDL-ESM2G, GFDL-ESM2M, HadGEM2-CC365, HadGEM2-ES365, in-mcm4, IPSL-CM5A-LR, IPSL-CM5A-MR, IPSL-CM5B-LR, MIROC-ESM-CHEM, MIROC5, MRI-CGCM3, and NorESM1-M. For map-based results, we report the median value across the years in each time frame and 19 models and for time series results we report the entire range of results across models and years. Where medians are calculated, we first calculate medians across years in each time frame and then calculate the median of these medians across the models. The RCP 4.5 scenario assumes a stabilization or reduction of greenhouse gas emissions starting around mid-century while the RCP 8.5 is an upper-end (90^th^ percentile) of many “no climate policy” scenarios that assume increasing emissions until the end of the century and is associated with relatively high temperature increases post mid-century compared with RCP 4.5 (van Vuuren et al., 2011). The temperature data were at a daily timestep and were then disaggregated into an hourly timestep using a sine-logarithmic algorithm (Linvill, 1990).

### 2.3 Dynamic Model

The Dynamic Model (Erez and Fishman, 1997; Erez et al., 1989), computes chill accumulation in units referred to as “chill portions” (CP). Chill accumulation is considered a two-step process. The first step is a reversible intermediary process where chill accumulates under low temperatures (a bell curve between -2°C and 13°C with optimal accumulation around 6°C), with no accumulation under -2°C, and potential for a negative accumulation by exposure to higher temperatures. The negation is a complex process that depends on the level, duration and cycle of exposure to high temperatures and only occurs during the intermediary process. Accumulation during this time can also be enhanced by exposure to moderately high temperatures (13°C-16°C) alternating with lower temperatures such as in a diurnal cycle. Once the intermediate accumulation threshold is surpassed, in the second step, chill accumulation increases by a unit called a chill portion unit. The accumulated chill portion units are conserved and cannot be reversed, and the intermediate level is set back to zero. Further details can be found in Erez and Fishman (1997); Erez et al. (1989).

The Dynamic Model implementation we used was provided as part of the chillR package for the R software (Luedeling, 2013)^1^

The function used is `chillR::Dynamic_Model(.)` and requires hourly temperature time series as input. The daily maximum and minimum temperature inputs from Section 2.2 were converted to hourly temperatures by the `chillR::stack_hourly_temps(.)` function in the chillR package that was based on Linvill’s work Linvill (1990). We used a simulation time frame of September 1^st^ to April 1^st^ covering the time frame for winter and spring phenology in our region. Chill accumulation in our area starts later than September 1^st^ and this start date ensures that the Dynamic Model can capture the inter-annual variation in the commencement of chill portions accumulation.

### 2.4 Chill Portion Requirements

Chill portion requirements are species- and cultivar-specific (Darbyshire et al., 2016; Luedeling et al., 2011). For the analysis to be applicable to a range of different tree fruit species and cultivars, we consider chill portion requirement thresholds from 20 to 75 at increments of 5 chill portions. This is based on the range of values compiled through literature reviews by Darbyshire et al. (2016) and Fruit & Nut Research & Information Center. Specific to varieties grown in the PNW, some of the known minimum chill portions requirements (Erez, 2000; Díez-Palet et al., 2019), are listed in Table 2 and they fall within the 20-75 chill portions range considered in this analysis.

**Table 2.**
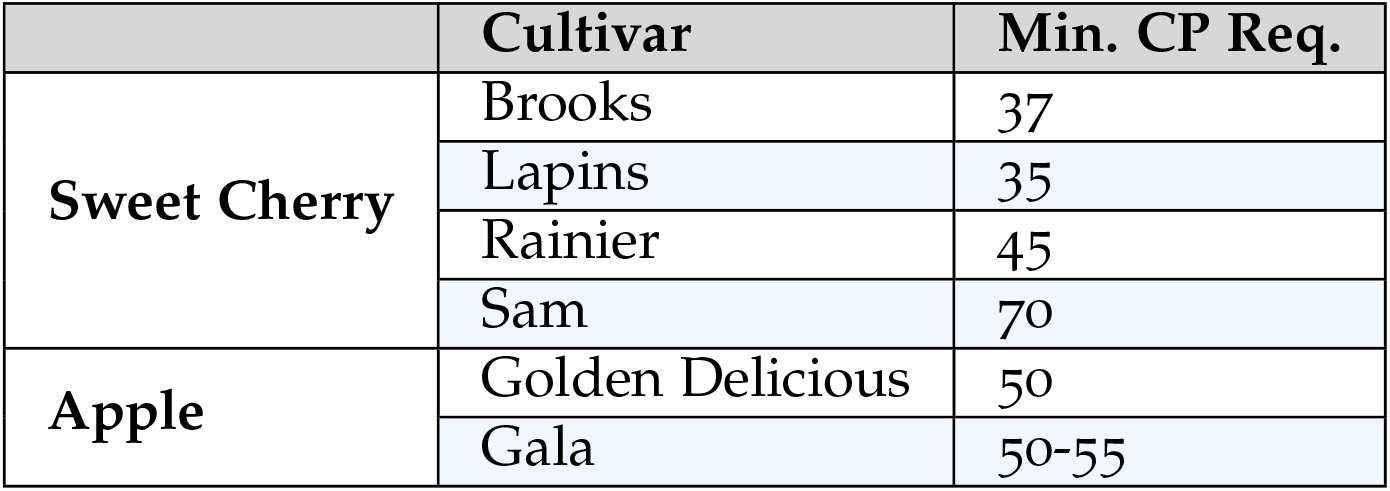
Chill Portions accumulation requirements for some commonly grown varieties in the PNW (Erez, 2000; Díez-Palet et al., 2019).

### 2.5 Heat Accumulation

In order to place shifts in chill accumulation within the context of potential shifts in heat accumulation, we use the growing degree day approach with a base temperature of 6.1°C, an upper threshold of 25.9°C, and the vertical cutoff approach. Growing degree day accumulation was calculated using the equations of Baskerville and Emin (1969), which fits a sine wave to the daily maximum and minimum temperatures, and calculates the area under the curve for heat unit accumulation. Historically, while temperatures amenable to heat accumulation do not commence until February or March in most of the PNW, we start accumulating growing degree days from December 1^st^ to allow to advancement in commencement of heat accumulation under warmer future climates.

## 3 Results and Discussion

In this section, regional spatial maps of median results are presented. Four representative locations (Fig. 1) are used to highlight temporal aspects and the spread of results from the 19 climate models.

### 3.1 Chill Portions Accumulation

#### 3.1.1 Magnitude of Chill Portions Accumulation

Historically, the southwestern tree fruit growing areas of Oregon State (Willamette Valley) accumulated the most, and the northern parts of Washington State accumulated the least chill portions (CP) units (Fig. 2). The apparent contradiction of the warmer areas (Willamette Valley) having the most CP units and the colder areas having the least is a function of colder areas having a higher exposure to temperatures lower than -2°C–temperatures that result in no CP accumulation. With climate change, modeled CP accumulation increased in the relatively colder northern parts of Washington state but decreased in other regions, with larger differences for the RCP 8.5 scenario. By the end of the century, in a switch from historical conditions, the northern parts of Washington state were projected to accumulate more CP than the southern regions in this study (Fig. 2).

**Fig. 2.**
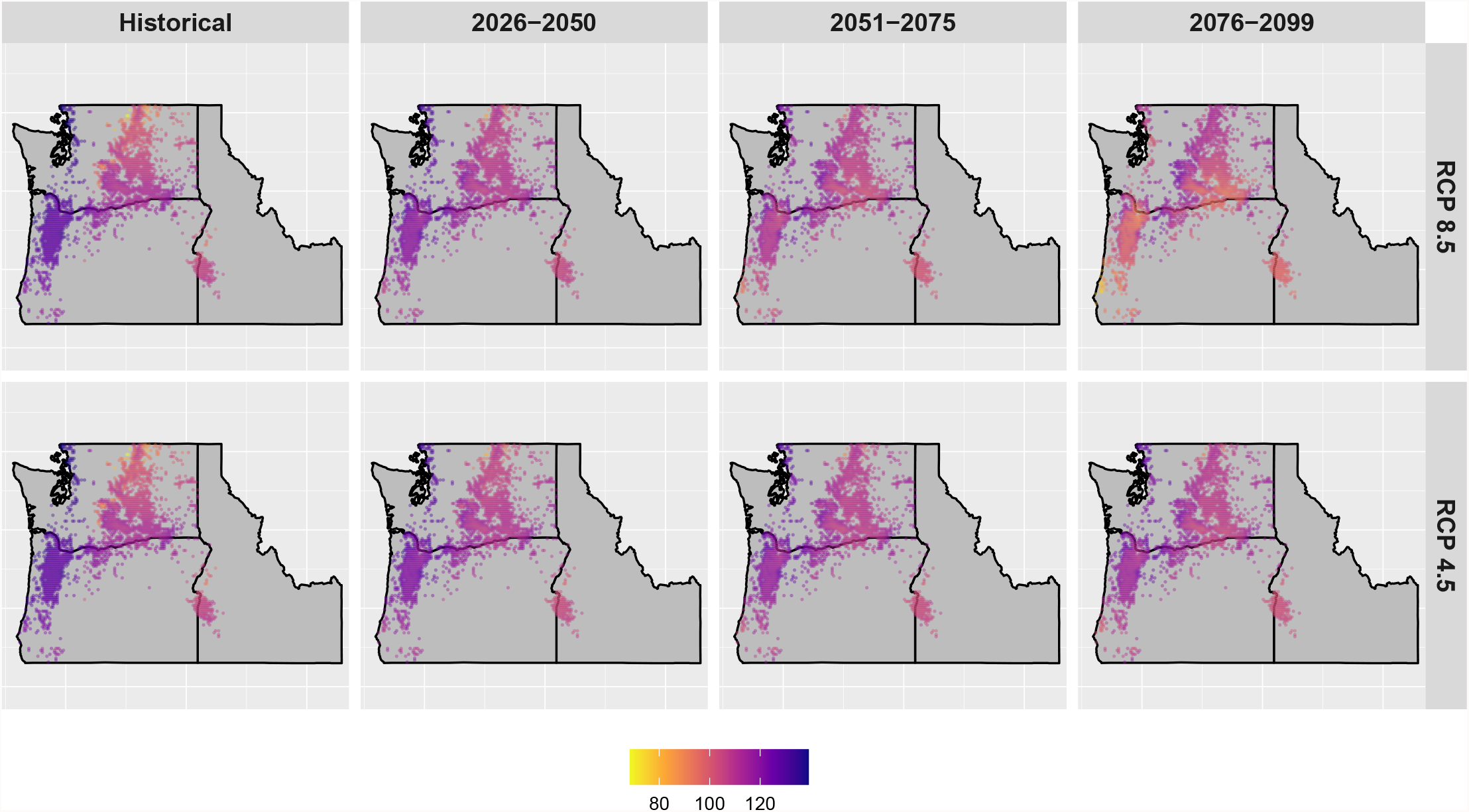
CP accumulation between September 1^st^ and April 1^st^ for different time frames under multiple climate models. First, the accumulated CP by April 1^st^ is computed for each location, year and model. Then for each location, the median of these values is taken over the years within each time window. Finally, the median of these median values is taken over the 19 climate models. Results are shown for the RCP 8.5 and 4.5 scenarios.

#### 3.1.2 Relative Change in CP Accumulation

Our simulations showed both increases and decreases across relatively short geographical ranges (Fig. 3). For the RCP 8.5 scenario, CP accumulation was generally projected to change by +/-25% for most areas with some coastal areas showing decreases larger than 40% (Fig. 3). The negative changes are smaller in magnitude for the RCP 4.5 scenario, especially after 2050, and positive changes are similar to that for RCP 8.5. However, for all regions, by the end of the century, an average of more than 80 CP were projected to accumulate by April 1^st^ (Fig. 2) for all 19 climate models, even under the RCP 8.5 scenario, implying that even with warming, projected temperatures are predicted to be conducive to chill accumulation. These levels of CP accumulation are above the range of minimum chilling requirements identified for different tree fruit species and cultivars in the PNW (Table 2). This suggests that there is limited risk of insufficient chill accumulation even for varieties with high chill requirements. This is in contrast to the results of Huston et al. (Houston et al., 2018) who quantified the risk for insufficient chill accumulation for two locations in the PNW; Corvallis (close to Eugene) and Wenatchee (close to Yakima). They noted that there is a likely risk of insufficient chill accumulation in Corvallis for high-chill-requirement varieties as early as the 2030s. The differences between our results and those reported by Huston et al. (Houston et al., 2018) are likely because they used the Chill Hour model while we used the Dynamic Model. The Chill Hour model does not account for dynamic conditions and studies using it tend to show exaggerated responses to warming compared to the Dynamic Model which is better able to capture changes to chilling conditions (Luedeling, Zhang, Luedeling and Girvetz, 2009; Luedeling, 2012). However, the general direction of change in Corvallis/Eugene is consistent across both studies. While Huston et al. (Houston et al., 2018) does not consider locations in northern Washington state, increases in chill accumulation in these colder regions are consistent with those reported by Luedeling et al. (Luedeling et al., 2011) in their global study which highlights changes in the Okanagan Valley in Canada which is just north of the Omak region in northern Washington state.

**Fig. 3.**
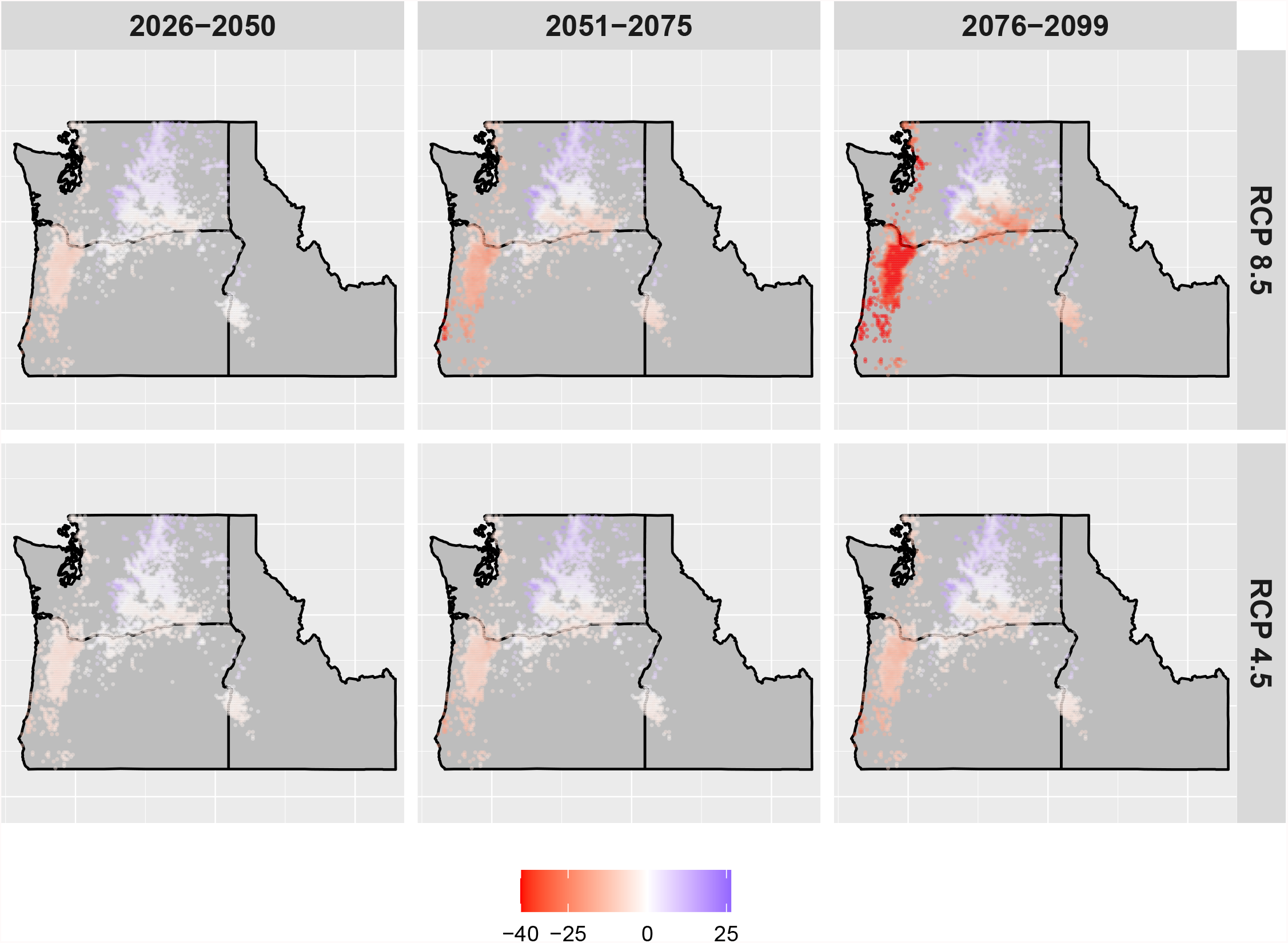
Percentage differences of CP accumulation between projections and historical observations. In this plot, for a given location and model, we computed the median of CP accumulation across years in three future time periods. Then, the differences between these projections and historical observations were computed. Then, we computed the median of percentage differences over the 19 climatic models. Chill seasons start on September 1^st^ and end on April 1^st^. Results are shown for the RCP 8.5 and 4.5 scenarios.

#### 3.1.3 Temporal Pattern of CP Accumulation and Spread Across Models

The temporal pattern of changes in CP as well as the spread across the different models for the four representatives locations shows the variable response (Fig. 4) in the rate of change over time and the different RCP scenarios.

**Fig. 4.**
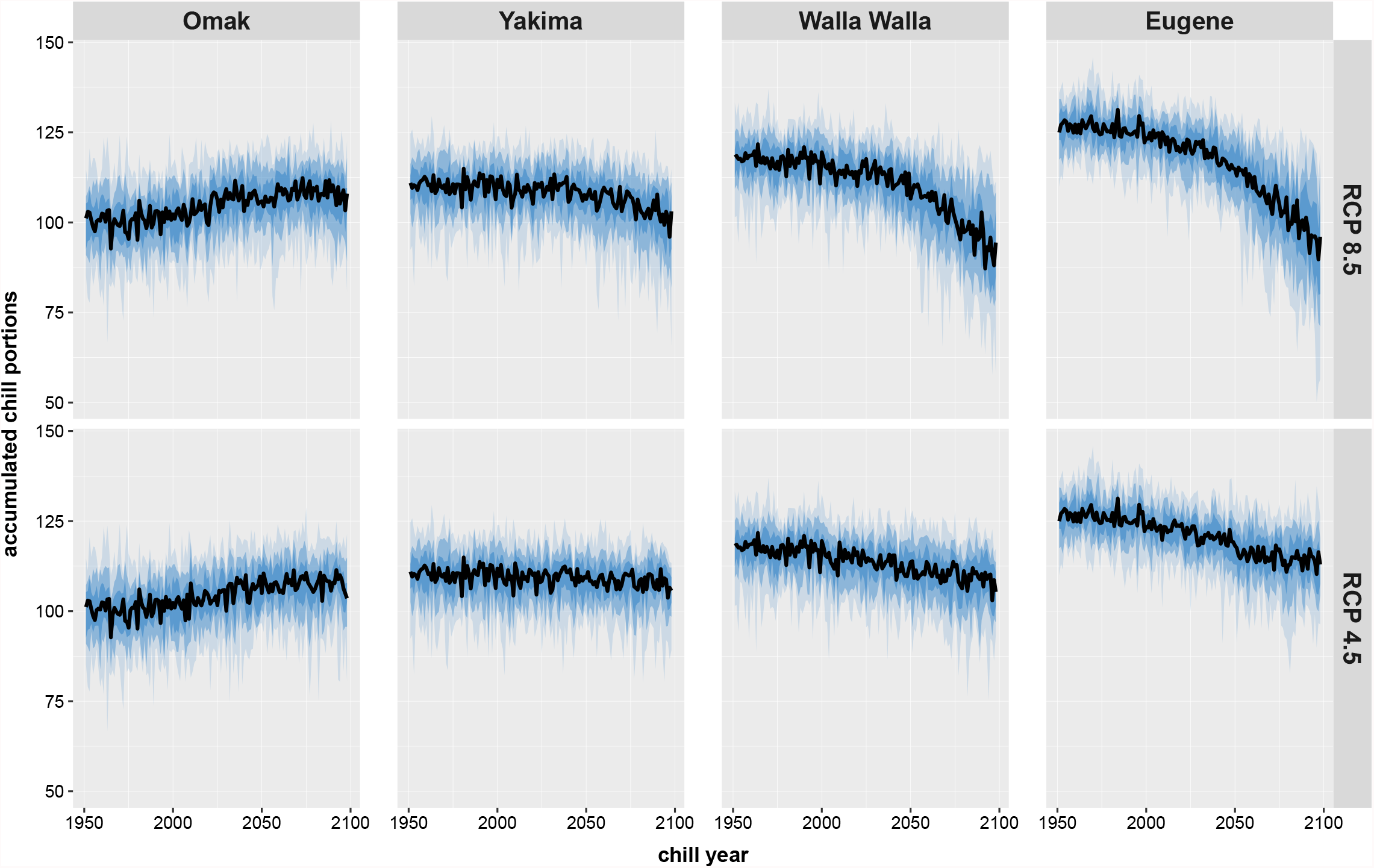
Temporal pattern of CP accumulation (September 1 - April 1) in Omak, Yakima, Walla Walla, and Eugene for the RCP 8.5 and 4.5 scenarios. The range for a given year corresponds to the 19 climate models. The darker shades covers the percentile range 25 to 75, the middle shade covers from 10^th^ percentile to the 90^th^, and the lightest shade goes from 0^th^ to 100^th^ percentile.

While Omak experienced increases in chill accumulation historically, the change seems to have stabilized and for RCP 8.5, the accumulated CP showed no change and remained relatively flat (Fig. 4). Under RCP 8.5, Yakima stayed flat until the mid-century and then it decreased at a nearly linear rate, while Walla Walla in southern Washington State and Eugene in Oregon State decreased at a nearly linear rate until the end of the century (Fig. 4). While there is a large spread across models, most models indicated meeting minimum CP requirements in all locations. There are just six climate model-year combinations that indicated a risk of not meeting minimum CP requirements for species and cultivars with high CP requirements (above 70 CP), especially in Oregon State (Eugene) and southeastern Washington State (Walla Walla) towards the end-of-century. The end-of-century minimum and maximum projected CP accumulation for Eugene (RCP 8.5) across the nineteen models were 56.4 and 119.2 CPs, respectively. For Omak, the minimum and maximum CP accumulations were 80.7 and 122.5 CP, respectively. Under RCP 4.5, changes are minimal as compared to RCP 8.5 as can be seen in Fig. 4.

#### 3.1.4 Reasons for Simulated Changes

To highlight the factors producing regional differences presented above, we examined the fraction of total hours (%) spent in different temperature ranges as relevant for the Dynamic Model (see Section 2.3, and references Erez and Fishman (1997); Erez et al. (1989)) for the simulation time frame of September 1–April 1 (Table 3). We focused on the RCP 8.5 scenario in this section as this is the scenario with relatively larger projected changes (Fig. 4). The increasing trend in CP accumulation for northern Washington State (Figs. 3 and 4) can be attributed to the models projecting significantly less time (−19%) with exposure to very cold temperatures (less than -2°C) which do not accumulate CP, and more time (7%) with exposure to a range of temperatures where CP accumulate. In contrast, other regions see decreases in the relative time of exposure to temperatures conducive to chill accumulation (−18% for Eugene). This, coupled with CP negating effects of temperatures outside the optimum range of -2°C to 13°C, contributed to more rapid decreases in CP accumulation for the warmer southern regions.

**Table 3.** The relative fraction of time (%) spent within specific temperature intervals between September 1^st^ and April 1^st^. These intervals correspond to differing effects on CP accumulation in the Dynamic Model. The results are for the RCP 8.5 scenario. These percentages correspond to hourly average temperature data aggregated across models and years.

| Location | Scenario | <-2°C | -2°C-13°C | 13°C-16°C | >16°C |
| --- | --- | --- | --- | --- | --- |
| Omak | Historical | 24.2 | 61.5 | 5.1 | 9.3 |
|  | RCP8.5 (2076-2099) | 4.9 | 68.8 | 7.3 | 19 |
|  | Difference | -19.3 | 7.3 | 2.3 | 9.7 |
| Yakima | Historical | 18 | 65.7 | 5.8 | 10.4 |
|  | RCP8.5 (2076-2099) | 4.5 | 65.2 | 9 | 21.2 |
|  | Difference | -13.5 | -0.5 | 3.2 | 10.9 |
| Walla Walla | Historical | 9.6 | 71.3 | 7.1 | 12 |
|  | RCP8.5 (2076-2099) | 1.7 | 61.3 | 11.8 | 25.3 |
|  | Difference | 7.9 | -9.9 | 4.6 | 13.3 |
| Eugene | Historical | 2.2 | 79.8 | 7.4 | 10.6 |
|  | RCP8.5 (2076-2099) | 0.3 | 62 | 14.4 | 23.3 |
|  | Difference | -1.9 | -17.9 | 7 | 12.7 |

### 3.2 Timing of Accumulation of Various Chill Portions

In general, most areas other than the northern region (e.g., Omak) see a projected delay in the timing for meeting CP thresholds (Fig. 5). For the RCP 8.5 scenario and the end of the century, models consistently projected that, compared with historical timings, the most southern location (Eugene) had a one-month delay in meeting CP thresholds and a little more than a 2-week delay for mid 21^st^ century projections. This delay was consistent with Houston et al. (2018). Although our magnitude of delay is lower than theirs, we are using different models. For the most northern location (Omak), models projected a delay in the timing for chilling fulfillment for the lower CP thresholds and an advancement for reaching greater CP thresholds with a reversal at thresholds around 45 CP in late December. This implies that increased exposure to temperatures optimal for chill accumulation is primarily in effect only after January for Omak. Even for central Washington State (Yakima), we can see that earlier delays are compensated later in the season with a shrinking delay. This, again, points to the wide variation in the response to warming within this relatively small PNW tree fruit production region.

**Fig. 5.**
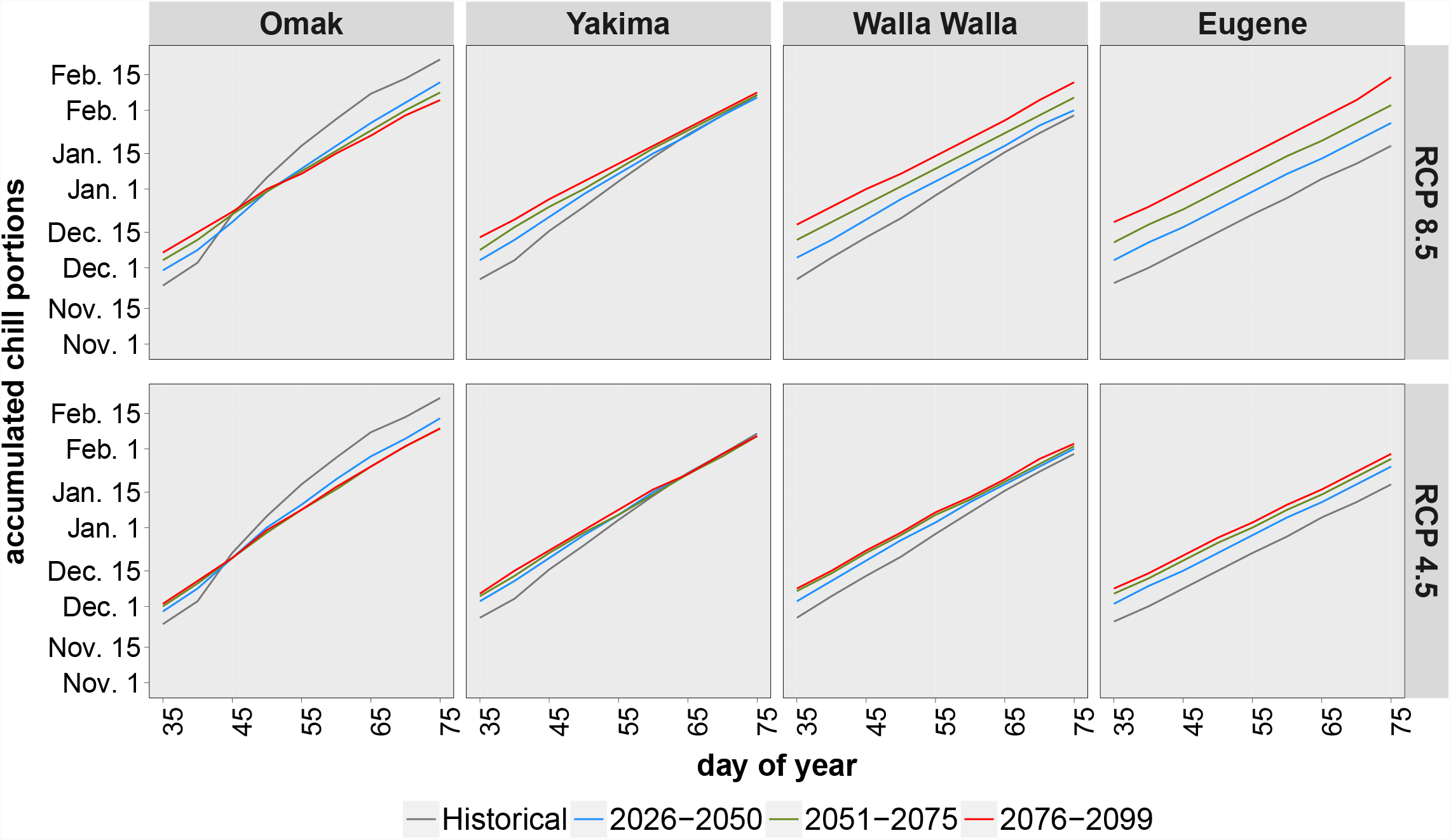
Day of the year (DoY) a range of CP accumulation threshold are met. These are medians of DoY values across years and across the 19 climate models.

Although the entire range of chill requirements (including the high values in the range) are met by April 1^st^, delayed fulfillment of chill accumulation thresholds have implications through affecting the overlap between chilling and heating processing and resulting in the dynamics between these processing becoming critical for spring phenology and bud break (Yang et al., 2020).

### 3.3 Overlap Between Chilling and Heat Accumulation Processes

In addition to delayed chill accumulation as noted in section 3.2, there are also advances in heat accumulation and are more pronounced for RCP 8.5 (Fig. 6) compared to RCP 4.5 (Fig. 7) suggesting an increasing future overlap between chilling and heat accumulation. In fact, historically, in most years, more than 75 chill portions were accumulated before heat accumulation started, indicating that, in these regions, spring phenology and bloom are primarily driven by forcing, consistent with the description of colder regions in (Yang et al., 2020; Guo et al., 2015). With warming, post-mid-century, there is potential for a regime shift away from being a region where spring phenology is primarily driven by forcing (and minimal or non-existent chill accumulation related production risks) to one where spring phenology is dynamically affected by both chill and heat accumulation processes, with a potential risk for issues such as non-uniform bloom and bud abscission that can result in production impacts (Atkinson et al., 2012). This may be only relevant for high-chill-requiring varieties since non-trivial chill portions continue to be accumulated well before the initiation of heat accumulation. The potential regime shift noted above indicated that simple thermal-time models may no longer be able to accurately quantify spring phenology, and other models which consider the dynamics between chilling and forcing requirements will be necessary. These models do not currently exist for the PNW. A better understanding of the dynamic interplay between chilling and forcing on spring phenology and the development of models that account for this in the context of PNW could become critical for future decision-support related to varieties with high chill requirements.

**Fig. 6.**
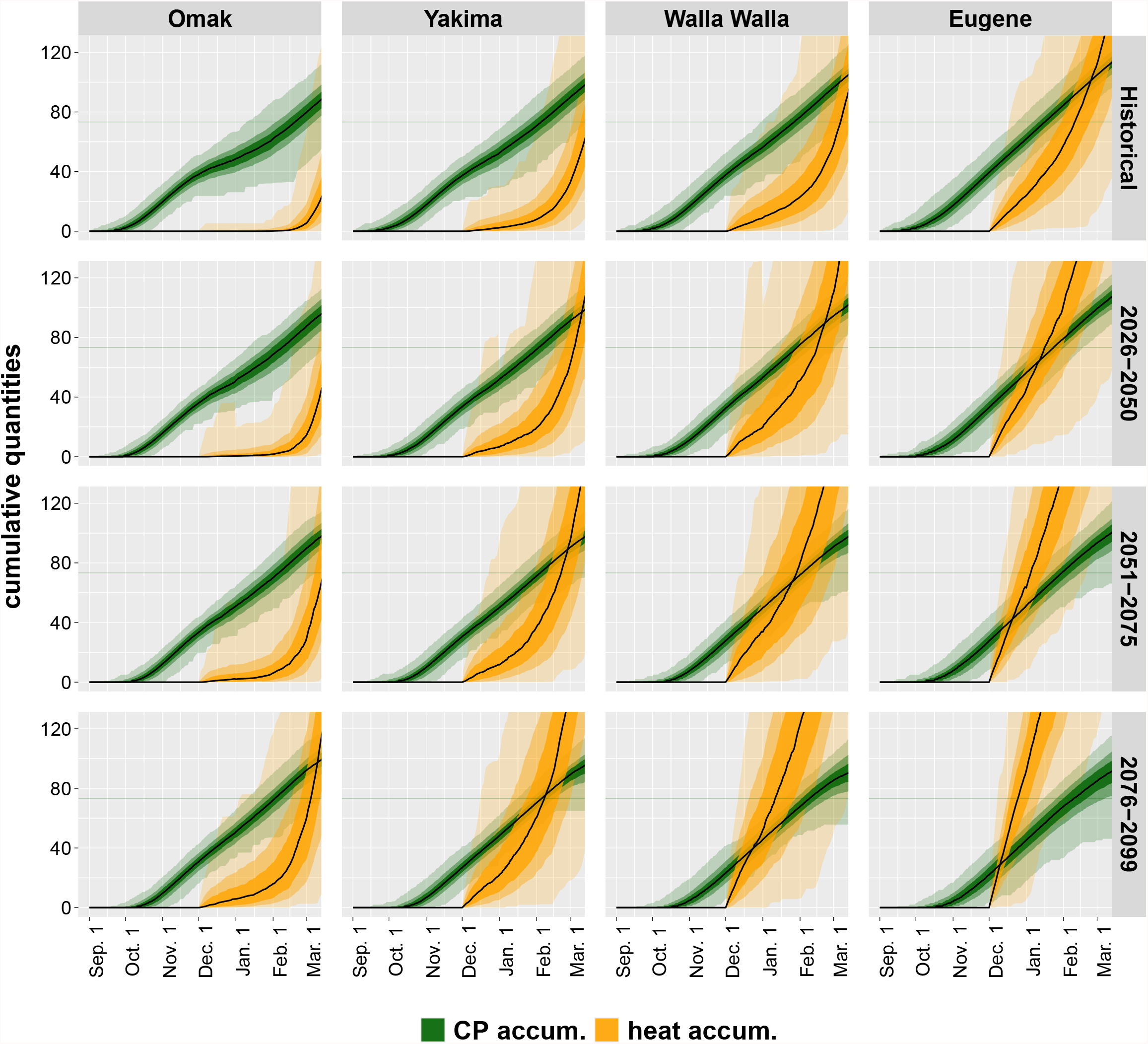
This plot corresponds to chill and heat accumulation for RCP 8.5. The green time series is of chill portions accumulation starting September 1^st^ and the orange heat accumulation in degree days starting December 1^st^. The range comes from 19 models and multiple years in the relevant time frame. The shadings correspond to 25^th^ to 75^th^ percentile (darkest shade), 10^th^ to 90^th^ percentile (lighter shade), and minimum to maximum (lightest shade) of CP accumulation. The middle black line is the median.

**Fig. 7.**
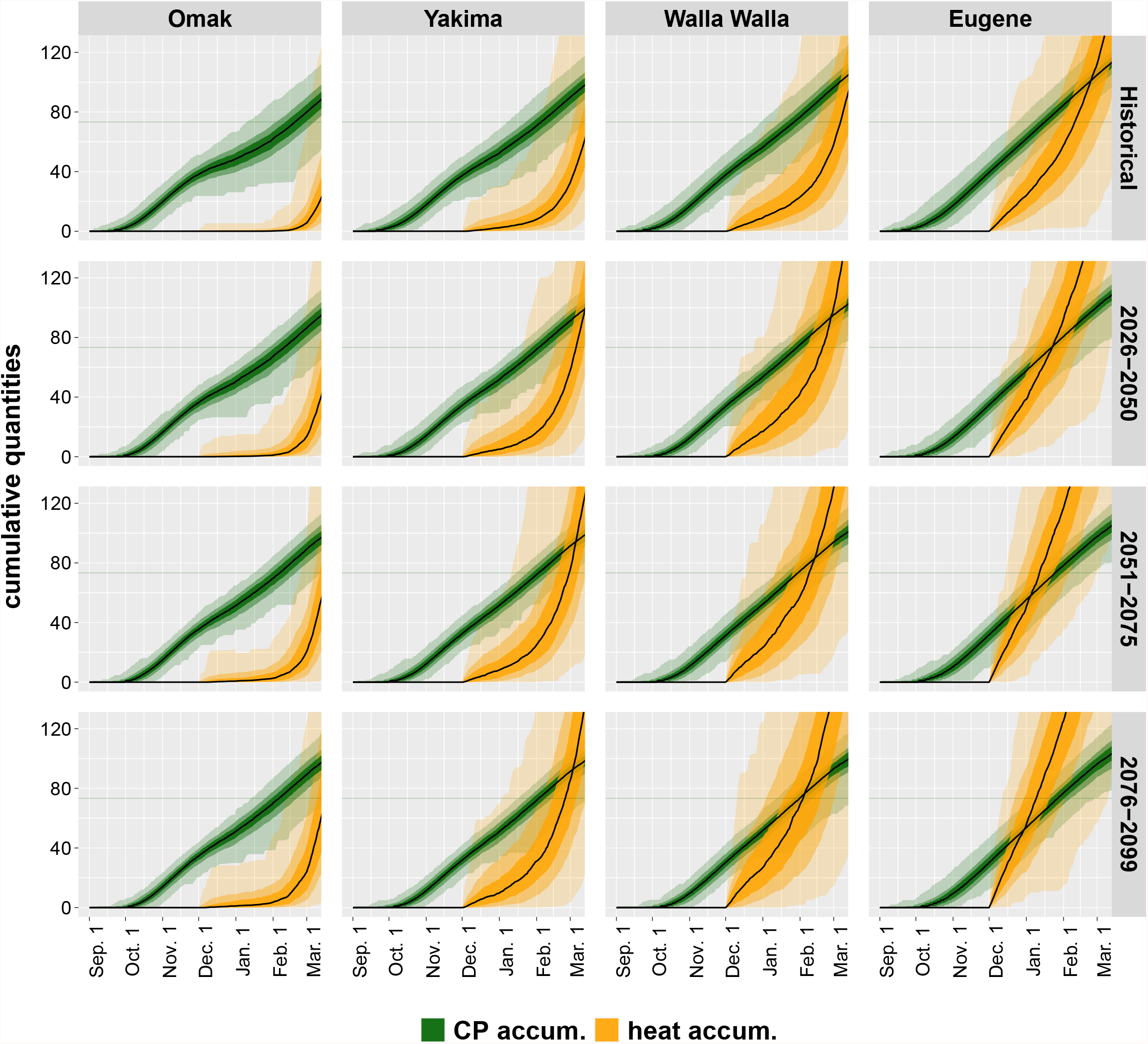
This figure is similar to Fig. 6 but the scenario is RCP 4.5 here. The green time series is of chill portions accumulation starting September 1^st^ and the orange heat accumulation in degree days starting December 1^st^. The range comes from 19 models and multiple years in the relevant time frame. The shadings correspond to 25^th^ to 75^th^ percentile (darkest shade), 10^th^ to 90^th^ percentile (lighter shade), and minimum to maximum (lightest shade) of CP accumulation. The middle black line is the median.

## 4 Conclusions

The tree fruit producing areas of the Pacific Northwest region of the United States experience large spatial variations in the changes in chill portions accumulation under climate change. The accumulation of chill portions under climate change is projected to decrease in more southern locations, stay the same in mid-locations, and increase in northern locations. The key factors contributing to spatial variation within the PNW can vary. Increased exposure to optimal chilling temperatures are projected to contribute to changes in the northern areas. In contrast, chill negation due to higher temperatures contribute to projected changes in other areas.

Even with warming, temperatures continue to be generally conducive to chill accumulation with no major expected issues with insufficient chill accumulation under most projections. However, a combination of delayed and reduced chill accumulation along with probable advancement in heat accumulation, might lead to many parts of the PNW transitioning from regimes where forcing is the primary driver for spring phenology to one where the dynamics and between chilling and forcing processes more strongly affects this phase change. This risk will likely manifest only in the high emission RCP 8.5, post-mid century, and for varieties with high CP requirements. Even though northern locations in Washington State face minimal risk, other warmer regions of the PNW with elevated future risk to chilling will likely continue as key production regions given their competitive advantage in terms of faster time to market. If the high emissions scenarios come to fruition, after 2050, these regions might need to plan for management strategies such as overhead irrigation for cooling, and chemical management of budbreak which can be costly interventions.

Significant gaps in our biological understanding of the winter dynamics and dormancy processes in tree fruit create uncertainty in quantifying the phenological response under climate change (Luedeling et al., 2011; Darbyshire et al., 2014; Richardson et al., 2013). Future modeling efforts must account for the overlap between fulfillment of chilling requirements (breaking of endodormancy) and environmental conditions promoting budbreak (break of ecodormancy), and the interactions between these two processes. It is under these conditions where risk to perennial tree fruit production exists and needs to be better understood, modeled, and projected. Given the significant spatial differences across a relatively small geographic range, it is also critical to understand, model and project these dynamics at a local landscape resolution.

## AuthorContributions

Conceptualization, H.N., K.R, L.K, M.J., and V.J.; methodology, H.N., L.K, K.R., V.J.; software, H.N; formal analysis, H.N.; resources, K.R.; data curation, H.N.; visualization, H.N.; supervision, K.R.; project administration, K.R.; funding acquisition, K.R., V.J., L.K.; writing—original draft preparation, K.R. and H.N.; writing—review and editing, V.J., L.K., M.J.

All authors have read and agreed to the published version of the manuscript.

## Acknowledgements

We thank Matthew Brousil for preliminary data processing.

## Data Availability

The datasets generated during and/or analysed during the current study are available from the corresponding author on reasonable request.

## Footnotes

1 Our code is available on Github (https://github.com/HNoorazar/Ag/tree/master/chilling).

